# Combined genetic and environmental factors shape the landscape of anti-fungal tolerance in *Nakaseomyces glabratus*

**DOI:** 10.1101/2025.07.02.662741

**Authors:** Sreyashi Acharjee, Edel M. Hyland

## Abstract

The rising incidence of fungal infections caused by *Candida* species is compounded by increasing antifungal resistance, compromising treatment efficacy. In *Candida albicans*, antifungal drug tolerance (AFDT) has emerged as a distinct, clinically relevant phenotype. However, AFDT remains underexplored in other clinically significant *Candida* species, and its prevalence across the genus is unclear. Here, we characterize AFDT in *Nakaseomyces glabratus* (formerly *Candida glabrata*), the second most common cause of candidiasis. Using disk diffusion and supra-MIC growth assays, we quantified AFDT across six clinical isolates and a reference strain.

Our results mirror findings in *C. albicans*, demonstrating reproducible AFDT across *N. glabratus* strains in response to multiple azoles. Importantly, tolerance did not correlate with resistance profiles, underscoring AFDT as a distinct phenotype. Isolates were categorized into high- medium- and low-tolerance groups based on variability. Highly tolerant strains showed reduced clearance in a *Galleria mellonella* infection model, aligning with previous AFDT studies. Tolerance was modulated by environmental factors such as media, temperature, and azole, yet relative strain tolerance remained stable—suggesting a heritable, strain-specific trait shaped by genetic and environmental factors. These findings underscore the clinical relevance of AFDT in *N. glabratus* and support further investigation into its molecular underpinnings and therapeutic impact.

## Introduction

Fungal pathogens affect over 250 million people worldwide and are responsible for approximately 1.5 million deaths annually, the rates of which are comparable to other widely known diseases such as malaria and tuberculosis (1). Furthermore, the diseases caused by fungal infections are of significant concern among immunocompromised individuals which include individuals infected with HIV, cancer patients, organ transplant recipients and among the elderly (2). These, *Candida* species which are the causative agents of candidiasis, account for nearly 75-90% of all fungal infections (3–6), with an associated mortality rate of upwards of 40-60% (7–11). Consequently, *Candida* infections represent a substantial public health concern(12, 13)

While *Candida albicans* remains the most frequently isolated species (1, 13), non-*C. albicans Candida* (NCAC) species, such as *Nakaseomyces glabratus* (formerly *Candida glabrata*), are emerging as significant threats (3, 8, 13, 14). *N. glabratus* is now recognized as the second most common cause of candidiasis (5, 15, 16), responsible for approximately 20–25% of *Candida* infections in the United Kingdom (<u>Source</u>). It is also a prominent cause of invasive fungal infections, particularly in individuals with weakened immune systems or impaired host defence mechanisms (16, 17)

Current treatment options for fungal infections are limited to just five classes of antifungal drugs, with azoles, polyenes, and echinocandins being the primary agents used in clinical settings (Reviewed in (18–21). Azoles—the focus of this study, target the cytochrome P450 enzyme lanosterol 14α-demethylase, encoded by the *ERG11* gene in yeast (18, 19). Inhibiting this enzyme disrupts the biosynthesis of ergosterol, the principal sterol component of the fungal cell membrane, thereby compromising membrane integrity and function (22). Azoles are generally fungistatic against yeasts, meaning they inhibit fungal growth without directly killing the cells.

A major factor limiting the efficacy of azole treatment is the rising prevalence of antifungal resistance (AFR) among pathogenic yeasts (23, 24). AFR is defined experimentally by an increase in the minimum inhibitory concentration (MIC) of a given antifungal agent. MICs are both drug and species-specific, and are determined as the lowest concentration of drug that visibly inhibits fungal growth in a microtiter-based assay (25). Currently, azole resistance rates in *C. albicans* isolates have been reported as ranging from 0-5 % depending on the geographical location (16, 26). However, for NCAC species like *N. glabratus*, this ranges from 2.6%–10.6%, even sometimes reaching up to 17% (8, 27, 28). Among these, *N. glabratus* presents a unique clinical challenge. Unlike other *Candida* species, *N. glabratus* demonstrates intrinsic reduced susceptibility to azoles, rendering it more difficult to treat effectively (10, 29). Moreover, *N. glabratus* has a heightened capacity to acquire resistance rapidly, as evidenced by the emergence of echinocandin resistance shortly after the drug class was introduced into clinical use (30, 31).

Although the genetic mechanisms of AFR have been well characterized, mutant strains do not fully account for the poor clinical outcomes observed in many *Candida* infections (32). A distinct phenomenon known as antifungal drug tolerance (AFDT) has recently emerged as an additional factor contributing to reduced drug efficacy (33, 34). Unlike AFR, which is driven by heritable genetic mutations, AFDT is primarily a phenotypic trait. Tolerant and non-tolerant cells within the same population are genetically identical, and the tolerant state is not stably inherited in the classical sense. Instead, AFDT is thought to arise from differential gene expression within a subpopulation of cells, leading to a transient, phenotypically heterogeneous response to antifungal stress (34).

The substantial body of evidence supporting antifungal drug tolerance (AFDT) is primarily derived from well-defined laboratory experiments (34). Tolerant cells are reproducibly detected in drug susceptibility tests such as disk diffusion assays and microbroth dilution assays when the typical 24 hour incubation time is extended to 48 hours. However, despite its recognition as an intrinsic characteristic of pathogenic yeast populations, drug tolerance remains poorly characterized, often going unnoticed and ignored in terms of treatment regimes. Indeed, clinical failures and recurrent infections can be attributed to a high tolerance phenotype rather than genetic resistance (33). Consequently, AFDT not only complicates treatment but also increases economic burden as it can often result in prolonged hospital stays (35). Of particular concern is the potential role AFDT plays in providing a window of opportunity for yeast populations to acquire true antifungal resistance mutations in the presence of antifungal drugs. (32, 33, 36).

To date, the study of AFDT mechanisms in *Candida* species has largely focused on *C. albicans*, leaving a gap in our understanding of how universal the phenotype is across other pathogenic yeasts. In this study, we focus on *N. glabratus*, given the clinical challenges associated with treating infections caused by this species. Using six *N. glabratus* isolates obtained from intensive care patients at Royal Victoria Hospital (Belfast), we observed that azole tolerance in *N. glabratus* shares many similarities with *C. albicans*. All strains maintained a subpopulation of tolerant cells and this did not correlate with the specific azole MIC for each strain. Additionally, we found that the specific level of tolerance was flexible, varying with environmental factors such as media, drug type, and growth temperature, yet this variation did not correlate with the strain’s doubling time under these conditions. What our data did suggest is that the primary determinant of AFDT in *N. glabratus* is genetic background, indicating a strong genetic component to this phenotype. Given the clinical implications of managing tolerant strains, understanding the underlying mechanisms driving this phenotype is imperative for improving treatment strategies of *N. glabratus* infections in the clinic.

## Results

### *N. glabratus* clinical isolates from Royal Victoria Hospital, Belfast are phenotypically diverse

The aim of this study was to observe and quantify antifungal drug tolerance in *N. glabratus* clinical isolates. We obtained six isolates (MLT031, MLT037, MLT040, MLT063, MLT071, and MLT080) from patients admitted to the intensive care unit at Royal Victoria Hospital, Belfast with systemic fungal infections. To assess the diversity of these isolates, we conducted standard phenotypic analyses in place of genetic sequencing. Each isolate was evaluated based on its doubling time and antifungal minimum inhibitory concentrations (MICs). As shown in Figure 1a, the reference strain ATCC2001, hereafter referred to as wild-type (WT), exhibited a doubling time (DT) of 1.5 hours in SC medium. All six clinical isolates demonstrated similar growth rates, with DT ranging from 1.4 to 1.6 hours. These findings suggest that, by this basic growth metric, the overall fitness of the clinical isolates is comparable to that of the WT strain.

**Figure 1.**
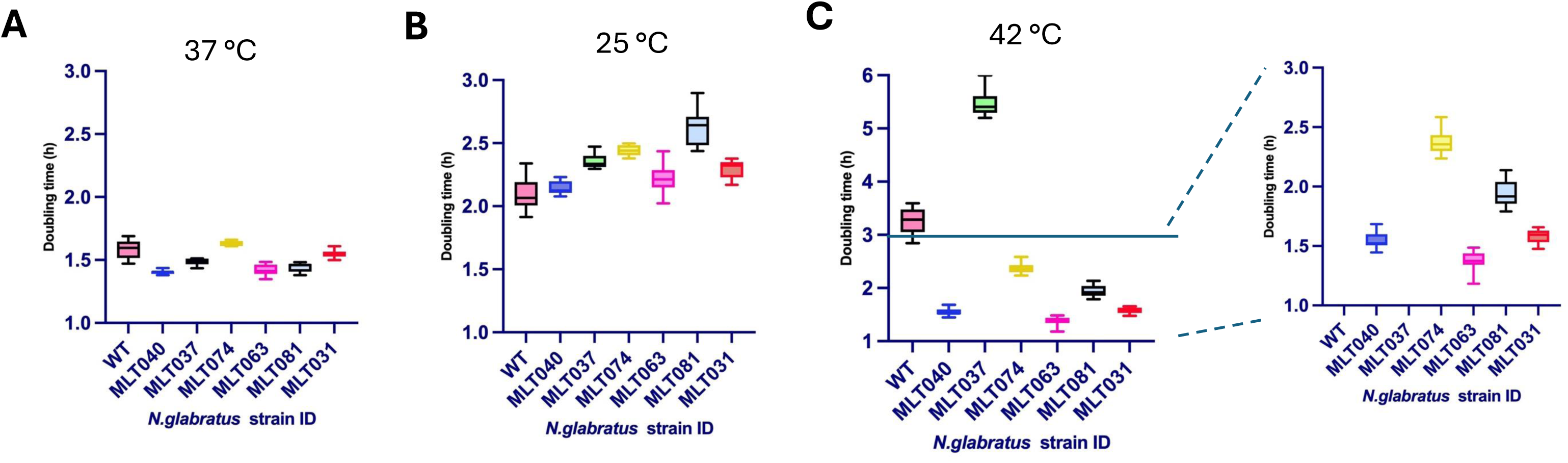
Variability in thermotolerance exhibited by *N. glabratus* clinical isolates. **(A-C)** represent the doubling time rates (hours) plotted against various clinical isolates of *N. glabratus* for temperatures 37 °C, 25 °C, and 42 °C. Data obtained by performing a microtitre plate growth assay in SC media and monitoring OD A600 over 24 hours. Data were analysed by GrowthCurver (44) and doubling times extracted. Data represent three biological replicates assayed in triplicate.

To assess the robustness of each isolate under environmental stress, we measured doubling times at sub-optimal temperatures of 25 °C and 42 °C. As shown in Figure 1b, responses to temperature shifts were variable across isolates. Several strains— including the wild-type (WT), MLT037, MLT071, and MLT081—exhibited significant changes in DT between temperatures, indicating temperature-sensitive growth. In contrast, isolates such as MLT031, MLT040, and MLT063 demonstrated more stable growth rates, suggesting greater thermal tolerance. This phenotypic stratification was most pronounced at 42 °C: while WT, MLT037, and MLT074 showed minimal or no growth, MLT031, MLT040, and MLT063 maintained detectable growth, with DT remaining below 100 minutes.

**Table 1:** Yeast strains analysed in this study.

| Strain | Species | Strain background |
| --- | --- | --- |
| ATCC 2001/CBS138 | *Nakaseomyces glabratus* | Wild type (WT) |
| MLT040 | *Nakaseomyces glabratus* | Clinical isolate from Royal Victoria Hospital, Belfast, ICU |
| MLT037 | *Nakaseomyces glabratus* | Clinical isolate from Royal Victoria Hospital, Belfast, ICU |
| MLT074 | *Nakaseomyces glabratus* | Clinical isolate from Royal Victoria Hospital, Belfast, ICU |
| MLT063 | *Nakaseomyces glabratus* | Clinical isolate from Royal Victoria Hospital, Belfast, ICU |
| MLT081 | *Nakaseomyces glabratus* | Clinical isolate from Royal Victoria Hospital, Belfast, ICU |
| MLT031 | *Nakaseomyces glabratus* | Clinical isolate from Royal Victoria Hospital, Belfast, ICU |

To evaluate antifungal drug resistance, we determined the MIC₅₀ values of all isolates against the azole drugs ketoconazole (KCZ), fluconazole (FCZ), and voriconazole (VCZ) (Figure 2, Table 2). VCZ and KCZ were selected due to their relatively lower MIC ranges for *N. glabratus* (34, 35). One isolate, MLT040, was more sensitive to FCZ than the wild-type (WT), and, along with MLT081, also exhibited increased sensitivity to both VCZ and KCZ. In contrast, MLT074 showed elevated resistance, with higher MIC₅₀ values for FCZ and VCZ compared to WT. MLT031 and MLT037 each displayed drug-specific resistance: MLT031 to FCZ and MLT037 to VCZ (Figures 2A and 2B). Interestingly, KCZ showed increased fungistatic activity in four of the six clinical isolates relative to WT (Figure 2C). As expected, MIC₅₀ values remained stable after 48 hours of incubation for all strain–drug combinations (Figure 2D). Together, these findings indicate that our clinical isolates represent a phenotypically diverse set, with variable responses to azole antifungals and temperature stress.

**Figure 2.**
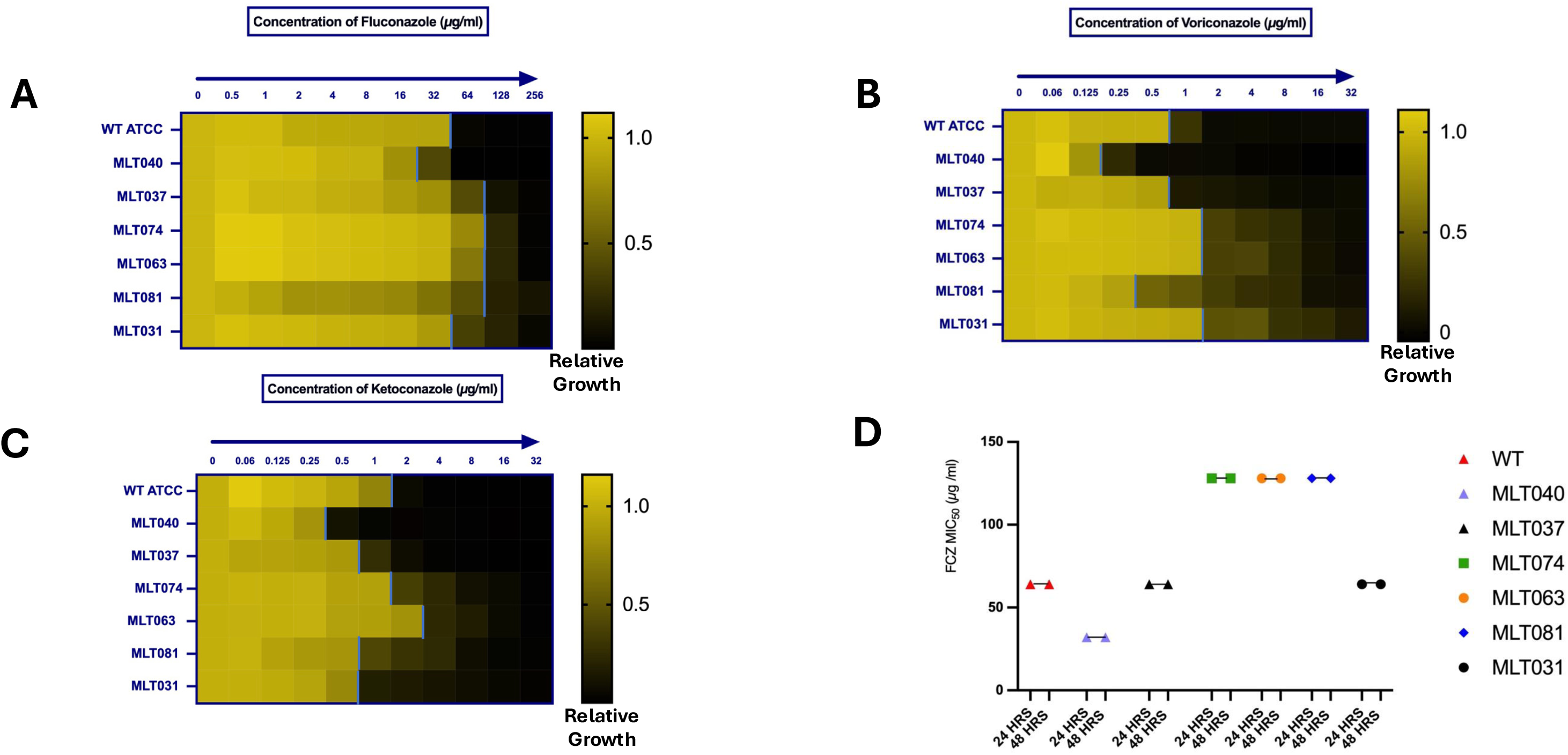
Clinical isolates of *N. glabratus* display distinct patterns of Fluconazole (FCZ), Voriconazole (VCZ), and Ketoconazole (KCZ) susceptibility. **(A-C)** MIC50 determination for all the tested clinical isolates referenced against the WT (typed ATCC2001 strain). The established CLSI protocol was implemented using 96 well plates, SC-complete media and 2-fold increases in concentrations of the indicated antifungal drugs. GrowthCurver analysis was undertaken on OD readings after 24 hours as in Figure 1. Data are visually represented using heat maps constructed using GraphPad Prism v10. vertical blue lines indicate the concentration at which growth is inhibited by >50%. **(D)** Represents the MIC50 comparison values between 24 hrs and 48 hrs time points, which remains constant across all the strains.

**Table 2:**
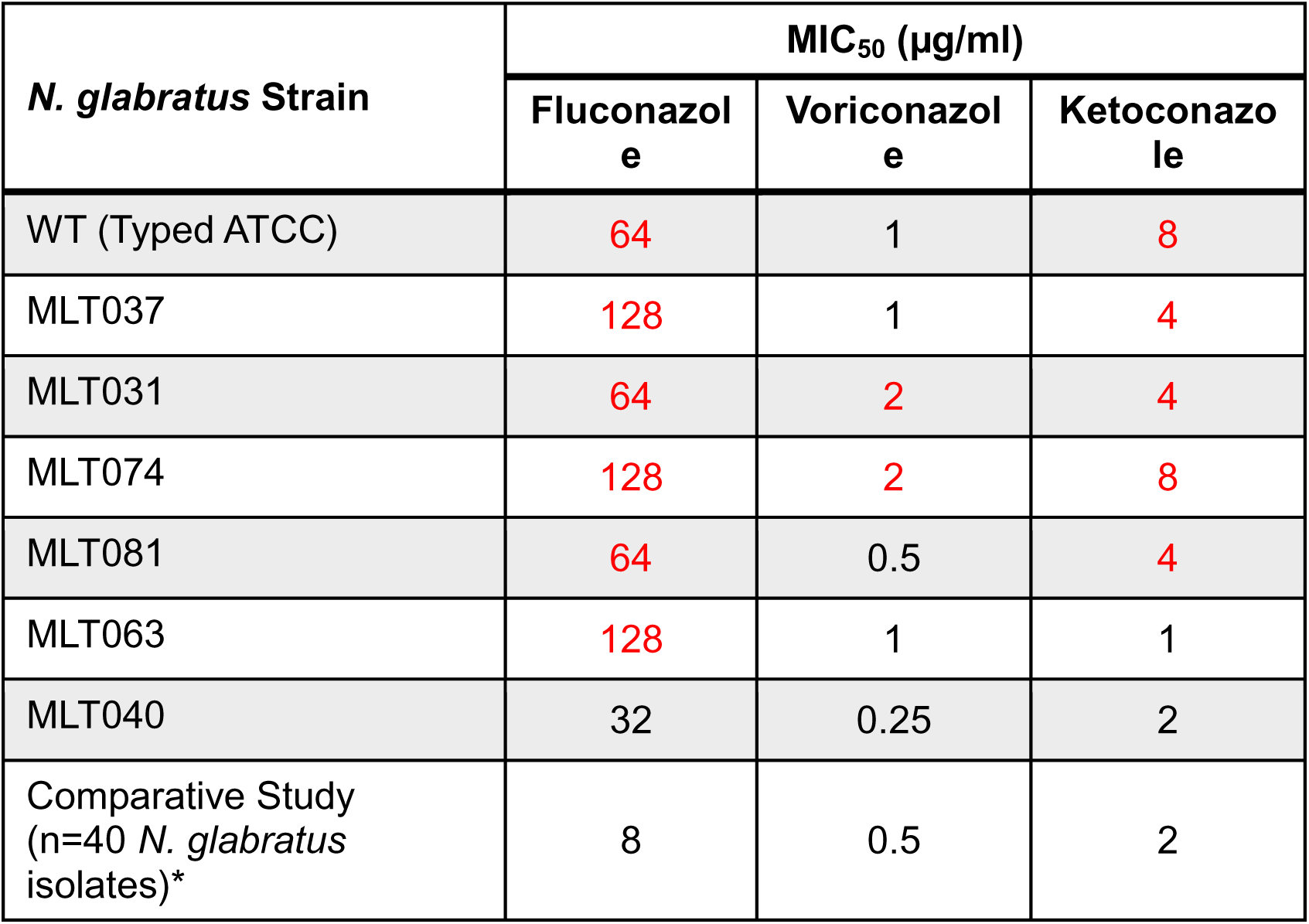
MIC_50_ (50% inhibition of growth) values for indicated strains, determined using CLSI methodology as described. Values in red denote increased resistance. *CSLI determined MIC_50_ (growth inhibited of 50% of isolates tested)(45)

### Clinical isolates of *N. glabratus* exhibit varying levels of fluconazole tolerance that are independent of their corresponding MIC₅₀ values

To assess and characterize the tolerance phenotype in our *N. glabratus* strains, we first quantified fluconazole (FCZ) tolerance under optimal growth conditions: YPD rich media and incubation at 37 °C (Figure 3). Using a disk diffusion assay (DDA) and calculating FoG₍₂₀₎ values, we classified the strains into three tolerance categories—high, moderate, and low. As shown in Figure 3A, strains MLT037, MLT031, MLT074, MLT063, and the wild-type (WT) were classified as moderately tolerant (FoG₍₂₀₎ range: 0.20–0.37), MLT081 as highly tolerant (FoG₍₂₀₎ > 0.58), and MLT040 as low tolerant (FoG₍₂₀₎ < 0.20). This classification was reproducible across three biological replicates (each tested in triplicate) and was not correlated with FCZ resistance. As shown in Figure 3B, a regression analysis between FoG₍₂₀₎ and RAD₍₂₀₎, a resistance metric derived from the same DDA after 24 hours (see Methods)—revealed a negative correlation (r = −0.583; R² = 0.33), supporting the distinction between tolerance and resistance phenotypes in *N. glabratus*. To further validate tolerance classification, we measured supra-MIC growth (SMG). Figure 3C presents both the raw and calculated SMG values for each strain, confirming the FoG₍₂₀₎-based categorization. Indeed, FoG₍₂₀₎ and SMG values were strongly correlated (r = 0.92; R² = 0.8587), as shown in Figure 3D.

**Figure 3.**
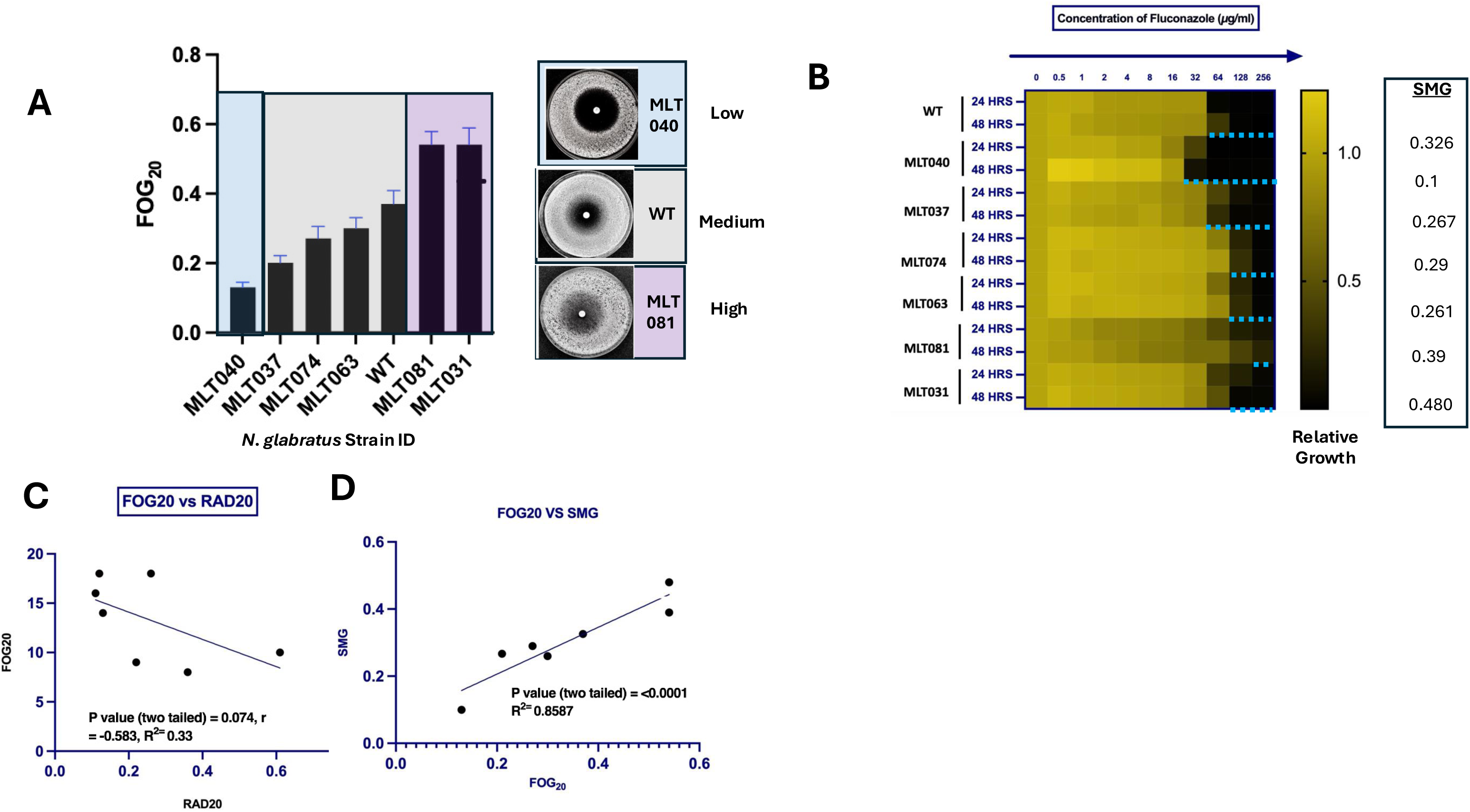
Clinical isolates of *N. glabratus* stratified into high, medium and low tolerant phenotypes. **(A)** FoG assays displaying fluconazole tolerance levels in indicated *N. glabratus* strains. Briefly 1 E+07 cells were plated on media plates, fluconazole (153 µg) disk was placed at the centre and plates incubated for 48 hours at 37 °C and photographed. Image was analysed and tolerance levels (high, moderate, low) were quantified using Disk Image R software. Error bars represent the standard deviation between three separate biological replicates assayed in triplicate. **(B)** Determination of Supra-MIC Growth (SMG) using microtiter-based growth assays in the presence of fluconazole. Growth was assessed by measuring optical density at 600 nm at both 24- and 48-hour time points. Horizontal dotted lines indicate the specific wells used for SMG calculation, as described in reference (25) **(C)** A negative correlation between RAD and FOG is observed in *N. glabratus* (r^2^ = 0.33), whereas in **(D)**, a positive correlation is seen between SMG and FOG (r^2^=0.86). Both SMG and FOG quantify drug tolerance, supporting the validity of these assays. Regression analyses were performed using the Pearson correlation coefficient test in GraphPad Prism v10.

### External factors modulate tolerance in *N. glabratus* clinical isolates

We next sought to determine whether tolerance levels in *N. glabratus* strains represent a stable phenotypic trait independent of external factors. The first variable we examined was the antifungal drug itself. Drug resistance mechanisms such as target site mutations are often drug-specific, even within the same class of antifungal agents (23). Therefore, acquired resistance to one azole does not necessarily confer cross-resistance to others. In contrast, since AFDT reflects a broader physiological response, we hypothesised that tolerance levels would remain constant across different azole drugs. To test this, we quantified AFDT to two additional azoles— voriconazole (VCZ) and ketoconazole (KCZ)—in all clinical isolates. As shown in Figure 4A, several strains exhibited statistically significant differences in tolerance depending on the drug. Specifically, strains MLT031, MLT081, MLT063, MLT074, and the wild-type (WT) showed significant differences in tolerance to VCZ or KCZ compared to fluconazole (FCZ) (p < 0.0001). For instance, MLT031 displayed high tolerance to VCZ but only intermediate tolerance to FCZ and KCZ. MLT037 also showed a small yet significant difference between FCZ and VCZ (p < 0.05), with greater tolerance to VCZ. In contrast, MLT040, classified as a low-tolerance strain, showed no variation in tolerance across the three drugs.

**Figure 4.**
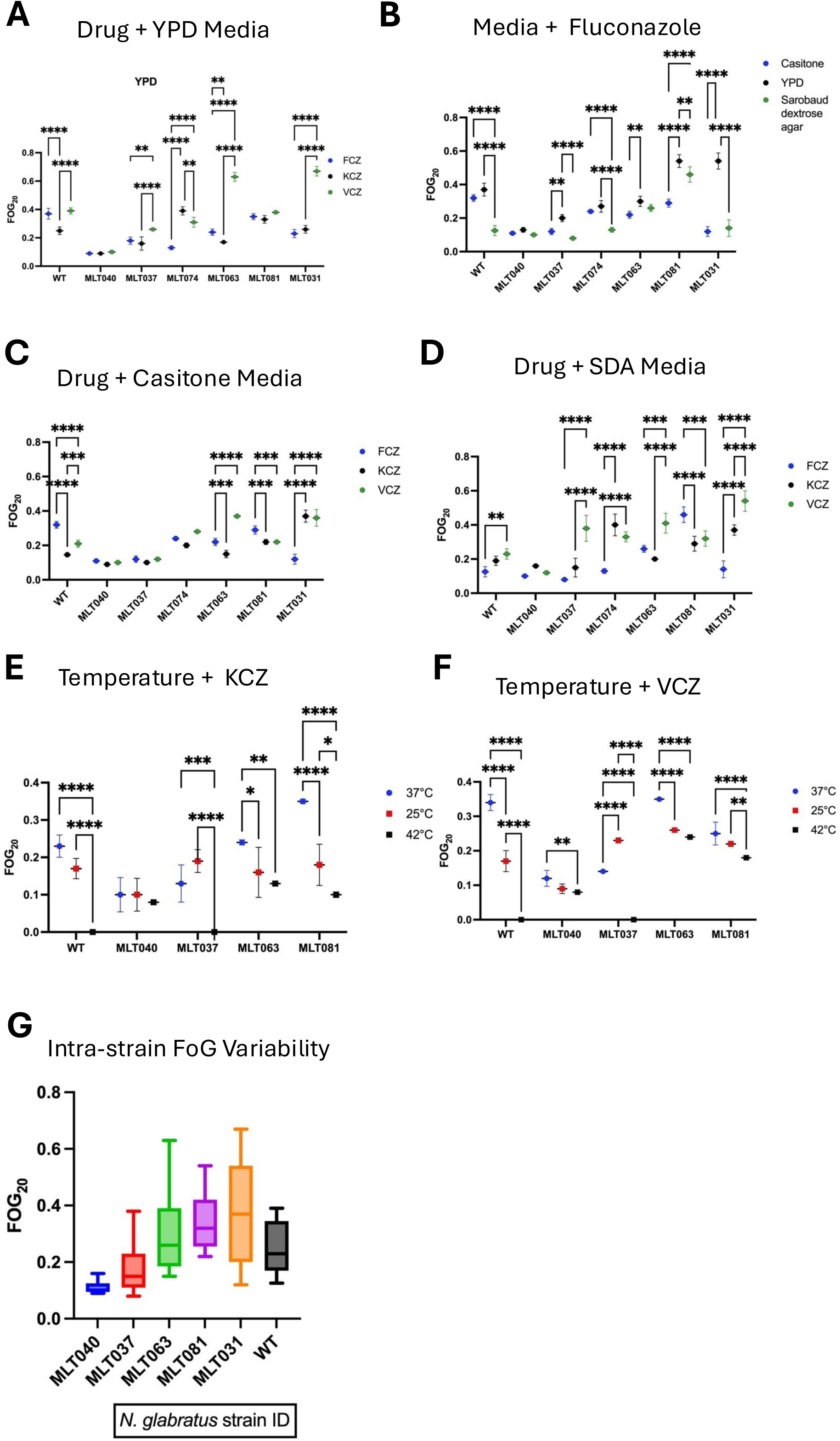
Tolerance levels in *N. glabratus* clinical isolates are determined by environmental cues: **(A-F)** FoG values of clinical isolates calculated as described in Figure 3 under the following conditions (A) YPD media and different azole drugs, FCZ=fluconazole, KCZ=ketoconazole, VCZ= voriconazole (B) Fluconazole and media variations, (C) Casitone minimal media and different drugs, (D) Sabouraud Dextrose Agar (SGA) and different drugs, (E) Ketoconazole, casitone media and varying temperatures, (F) Voriconazole, casitone media and varying temperature. Error bars represent the standard deviation between three separate biological replicates. The statistical tests performed are determined by a two-way Annova and the p-value ranging between (<0.0001-0.005) with a significance level ranging from highest (****) to lowest (**). (G) All FoG_20_ values (in panels A-F) for a given isolate was plotted to visualise the intra-strain variability. This illustrates the impact of environmental variables on tolerance levels for each of the strains.

Next, we assessed the influence of media composition on tolerance to individual azoles, with a specific focus on the role of nutrient availability. We hypothesized that nutrient limitation might enhance tolerance through two possible mechanisms: (1) slower growth rates could reduce the need for membrane biosynthesis, thereby lowering sensitivity to azoles, and (2) nutrient deprivation might activate stress response pathways, resulting in a more resilient, tolerant subpopulation (28). To test this, we repeated our tolerance assays using casitone medium, which lacks specific minerals and trace elements and Sabouraud Dextrose Agar (SDA), a nutrient-rich medium with a lower pH than YPD. In total, we conducted approximately 60 tolerance assays across all pairwise combinations of drug and media, with each condition tested in at least triplicate (Figures 4B–D).

As shown in Figure 4B, six of the seven strains exhibited statistically significant changes in tolerance across different media (p < 0.01–0.0001). Our results reveal a complex picture. For one, all strains showed poor growth on SDA (data not shown), and fluconazole tolerance in the moderately tolerant strains (MLT037, MLT074, MLT063, and WT) was lower on SDA compared to other media (Figure 4B). However, tolerance to other drugs was not similarly reduced in these strains on SDA—in fact, voriconazole tolerance was higher on this medium. This suggests that the observed phenotype results from an interaction between the drug and the specific media conditions for these strains.

Secondly, tolerance measured on casitone medium was the most consistent and reproducible across all strains and drug combinations. As shown in Figure 4C, 4 out of 7 strains tested displayed highly stable FoG_20_ values. In contrast, the high tolerant strains MLT081 and MLT031 showed the most variability in our assays, indicating that their high-tolerance phenotype is strongly dependent on both the drug and media composition. For example, MLT031 exhibited FoG_20_ values on SDA ranging from 0.14 with fluconazole to 0.54 with voriconazole—classifying it as both low and high tolerant under different conditions (Figure 4D). For MLT081, high tolerance was primarily observed in the standard YPD + fluconazole condition. On casitone medium (Figure 4C), MLT081 tolerance levels were reduced across all drugs, and on SDA, tolerance decreased further for both ketoconazole and voriconazole. These results suggest that media composition has the greatest influence on drug tolerance in MLT081.

The final variable we assessed was temperature, using DDA conducted at 25 °C and 42 °C. As illustrated in Figures 4E and 4F, maximal tolerance was observed at 37 °C—the optimal growth temperature for *N. glabratus*—in five out of six strains. The one exception was MLT037, which exhibited increased tolerance to all azoles at 25 °C.

As previously shown in Figure 1, thermotolerance varied among the isolates. Both the wild-type (WT) strain and MLT037 failed to grow at 42 °C, while all other strains displayed minimal growth impairment at this elevated temperature. Importantly, the trends observed in tolerance across media composition, azole type, and temperature were fully supported by results from the supra-MIC growth (SMG) assay (not shown) reinforcing the robustness of the observed tolerance phenotypes across experimental conditions.

To facilitate interpretation of the dataset, we next evaluated the phenotypic stability of drug tolerance across all tested conditions for each strain. As illustrated in Figure 4G, distinct patterns of variability were observed. The low-tolerance strain MLT040 exhibited minimal fluctuation in FoG_20_ values, indicating a highly stable tolerance phenotype. In contrast, MLT031 demonstrated pronounced variability in FoG_20_ measurements across conditions, consistent with an unstable tolerance phenotype. The remaining three strains, including WT, showed moderate and relatively stable tolerance, with FoG_20_ values generally ranging from 0.2 to 0.4. However, in very specific conditions, these values did increase, highlighting the influence of environmental factors on the expression of this phenotype.

### Azole drug tolerance is not defined by growth rate in *N. glabratus*

Based on the results thus far, we conclude that azole tolerance in *N. glabratus* is not a fixed trait but rather responds dynamically to external factors such as temperature, nutrient availability, and drug exposure. However, the observed responses did not follow a uniform trend; the directionality of change in response to each stimulus was predominantly strain-specific.

To interpret these findings, we hypothesized that tolerance levels might be negatively correlated with growth rate of a strain in a given set of conditions. To test this, we plotted the quantitated doubling time (DT) values from our SMG growth assays for each condition against the corresponding FoG₍₂₀₎ values obtained from the DDA assays. Our results, shown in Figure 5, demonstrate a negligible or weak correlation between growth rate and tolerance level, indicating that the key biological factor defining AFDT in *N. glabratus* is *not* the rate of cell division.

**Figure 5.**
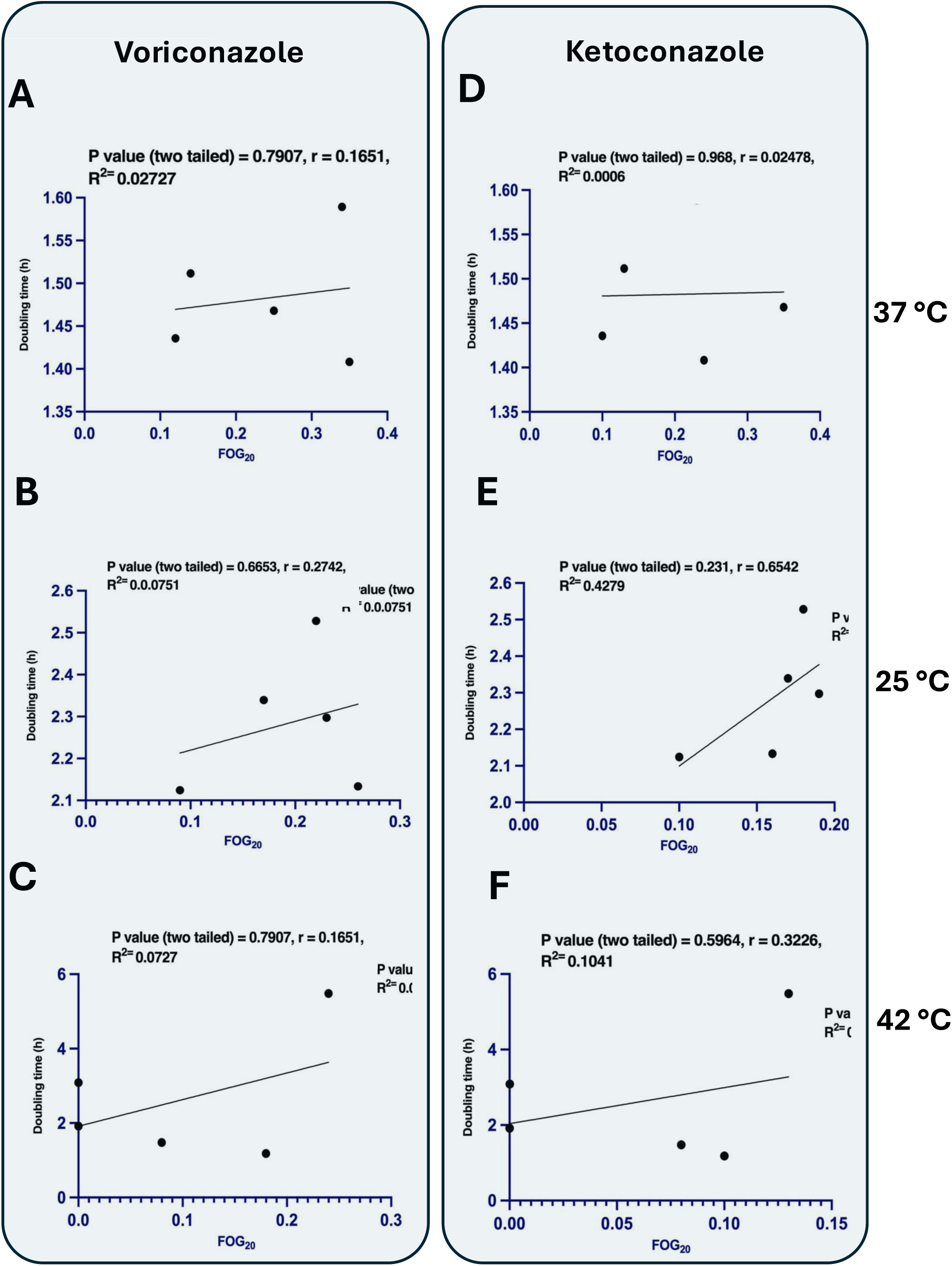
Non-significant correlation between tolerance levels and growth rate for *N. glabratus* isolates. Data are plotted as doubling time (hours) against FoG_20_ values for the 4 clinical isolates, MLY040, MLT063, MLT081, MLT031, and WT. Doubling times were calculated in the absence of drug at the indicated temperatures, 37 °C, 25 °C and 42 °C. FoG_20_ values obtained using DDA as in figure 3 with **(A-C)** Voriconazole (1µg) treatment or **(D-F)** Ketoconazole (50 µg) treatment. Data represent mean of biological triplicates.

High-tolerance *N. glabratus* strains exhibit reduced responsiveness to azole treatment *in vivo*.

High-tolerance *C. albicans* strains have been shown to reduce the efficacy of fluconazole treatment in both patients and murine models of candidiasis (37). To evaluate whether a similar trend is observed for *N. glabratus*, we monitored the effect of FCZ treatment of *G. mellonella* infected with high (MLT081) and low (MLT040) tolerant strains. Initially, we scored worm survival over 7 days for each infection, with or without antifungal treatment. As shown in Figure 6A, the trends in % survival aligned with the observations in *C. albicans*. MLT081 (high tolerance) was the most virulent strain, with only a 10% survival rate after 7 days, which increased to 25% upon treatment with FCZ. In contrast, MLT040 (low tolerance) had a 20% survival after 7 days, which rose to 40% in the presence of FCZ. However, none of these differences were statistically significant. Therefore, we sought another metric to monitor the physiological impact of high and low tolerant strains and chose to monitor the formation of cocoons in each treatment group over 7 days. Cocooning rate is a measure of the overall fitness of the worm whereby the higher the rate, the healthier the worm (38). Figure 6B indicates that > 50% of the individuals in the uninfected control groups had formed cocoons by day 7, whereas those infected with *N. glabratus* regardless of treatment, were significantly less able to develop into cocoons, consistent with compromised hosts. However, yeast strain tolerance level did not differentially impact cocoon formation, suggesting that AFDT does not influence the progression of an *in vivo* infection.

**Figure 6.**
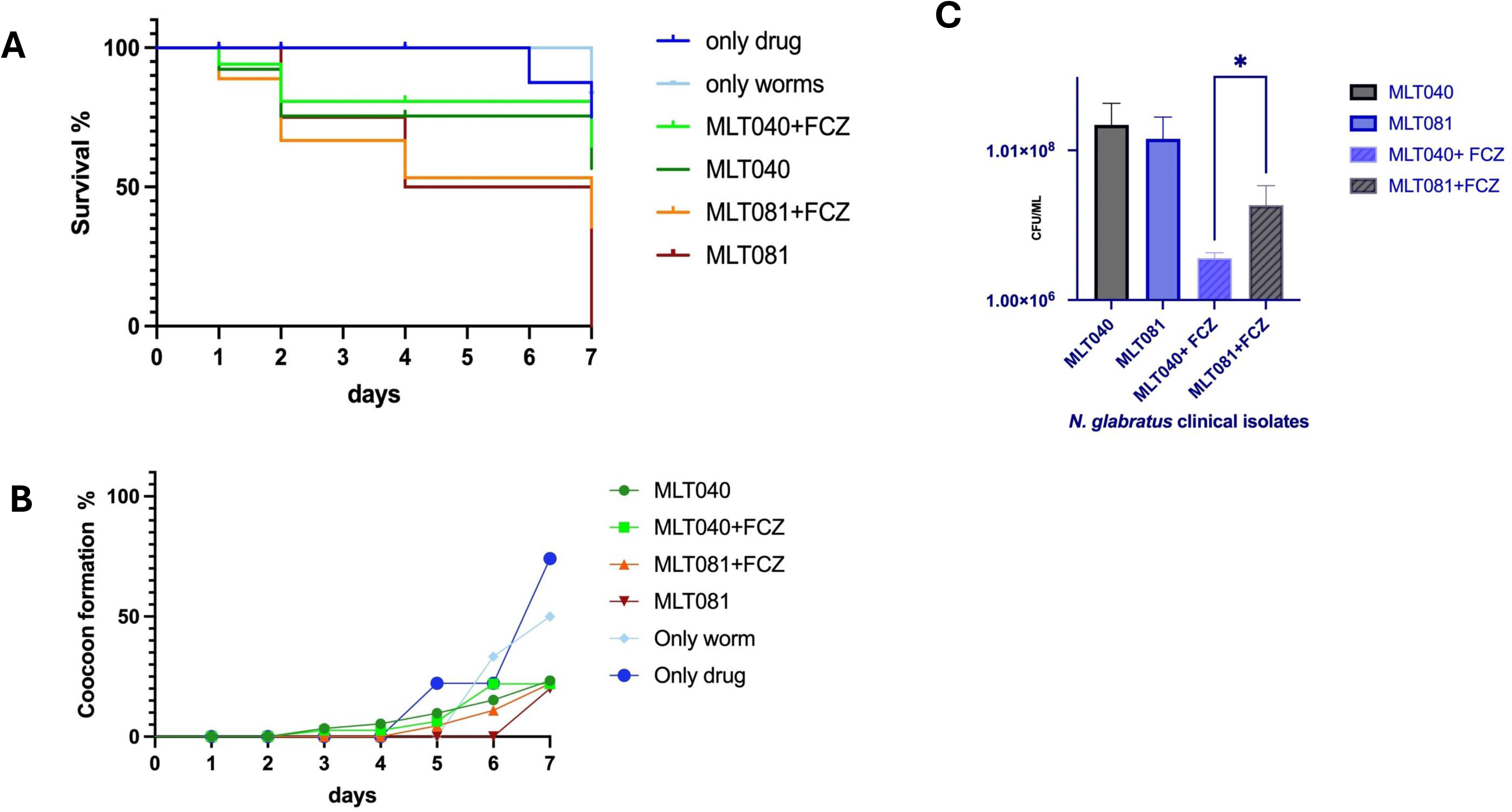
Efficacy of fluconazole treatment is reduced in *G. mellonella* infected with high tolerant *N. glabratus isolates*. **(A)** Seven-day survival curve of *G. mellonella* infected (n=20) with 5 E+06 *N. glabratus* cells with or without fluconazole treatment (12 mg/kg, or 4.2 µg total per worm). MLT040 = low tolerant strains, MLT081 = high tolerant strain. **(B)** Cocoon formation of *G. mellonella* throughout 7 days *N. glabratus* infection as in (A). **(C)** Fungal load (cfu/ml) of indicated experimental conditions. Briefly, at day 7 post infection, haemocoel from the last-pro leg of surviving worms (n=3) were collected and plated at various dilutions on YPD rich media containing chloramphenicol (100 ug/ml). Following 24 hours incubation 37 °C, colonies were counted.

That said, we did uncover a significant difference in treatment efficacy between high and low-tolerant strains. On day 7 of infection, we extracted the worm haemocoel and calculated fungal load in terms of CFU/ml. Figure 6C shows that although fungal load was comparable between high (MLT081) and low (MLT040) tolerant strains in untreated infections, there was a significant difference upon identical FCZ treatment. MLT040 fungal load was reduced by 4.1-fold, whereas MLT081 was more difficult to treat, with a reduction in CFU/ml corresponding to only 1.7-fold. Furthermore, two-way annova tests revealed a significant difference of p < 0.05 (*) between the high (MLT081) and the low-tolerant (MLT040) strains in response to treatment with fluconazole.

## Discussion

The emergence of antifungal drug tolerance (AFDT) as a clinically significant phenotype in fungal pathogens has challenged the traditional dichotomy between susceptible and resistant classifications. While extensively characterized in *Candida albicans*, the extent to which AFDT contributes to drug failure in non-*albicans* species has remained largely unexplored. In this study, we provide one of the first detailed characterization of azole tolerance in *Nakaseomyces glabratus*, the second most common cause of candidiasis, and demonstrate that AFDT is a robust, quantifiable, and strain-specific trait in this species.

Our results reveal that clinical isolates of *N. glabratus* exhibit reproducible AFDT that is independent of antifungal resistance. This distinction reinforces previous work in *C. albicans*, where AFDT has been shown to arise from phenotypic heterogeneity rather than heritable resistance mutations (32, 33). Indeed, both disk diffusion (FoG₍₂₀₎) and broth microdilution (SMG) assays produced concordant tolerance classifications, confirming the reliability and reproducibility of tolerance metrics.

Importantly, our data also demonstrate that AFDT is modulated by external factors including temperature, media composition, and azole type. These findings echo the dynamic nature of tolerance observed in other microbial systems, where environmental stressors can trigger physiological adaptations that promote drug survival. For example, we observed that nutrient-limiting media such as casitone could either enhance or suppress tolerance depending on the strain, suggesting the activation of strain-specific stress responses.

Similarly, tolerance fluctuated with temperature, peaking at the optimal growth temperature (37 °C) for most isolates but increasing at 25 °C in MLT037 suggesting that AFDT can be influenced by strain-specific thermotolerance mechanisms. This observation is supported by studies in *C. albicans* which demonstrated that isolates can express either temperature-enhanced tolerance (TET), or all temperature tolerance (ATT). (39)

Nevertheless, despite this plasticity, tolerance rankings were largely preserved across conditions, indicating that relative AFDT is a stable, strain-specific trait likely underpinned by genetic differences. This was further supported by the observation that tolerance was not significantly correlated with growth rate, suggesting that AFDT is not simply a function of metabolic activity or proliferation speed, but instead represents a distinct physiological state or regulatory program. The lack of a consistent link between growth rate and drug tolerance in *Candida* species is well-documented in scientific literature (33, 34, 40). This dissociation underscores the complexity of antifungal tolerance mechanisms, which often operate independently of traditional growth metrics.

The *in vivo* relevance of these findings was further supported by our *Galleria mellonella* infection model. While overall virulence (as measured by survival and cocooning) did not significantly differ between high and low-tolerance strains, antifungal treatment outcomes did. Fluconazole was significantly less effective at reducing fungal burden in larvae infected with the high-tolerance strain compared to the low-tolerance strain. These data parallel findings in *C. albicans* and provide compelling evidence that AFDT directly undermines antifungal efficacy *in vivo*, even in the absence of resistance—supporting its clinical relevance.

Taken together, our findings establish AFDT as a prominent, heritable, and therapeutically significant phenotype in *N. glabratus*. The presence of a persistent subpopulation of tolerant cells may not only contribute to treatment failure but could also promote the emergence of true resistance by allowing survival under sustained drug pressure (30, 36). Moreover, the diversity in tolerance observed among clinical isolates underscores the need to incorporate tolerance assessments into standard antifungal susceptibility testing, especially for difficult-to-treat NCAC species.

Future investigations should prioritize elucidating the molecular determinants underlying AFDT in *N. glabratus*, with particular emphasis on regulators of stress response pathways, drug efflux mechanisms, and epigenetic modulators. Integral to these studies is the implementation of single-cell analyses within clonal populations to resolve cell-to-cell variability in transcriptional landscapes, intracellular drug kinetics, and chromatin architecture. Such high-resolution approaches will be essential for defining population heterogeneity and mapping the distribution of AFDT phenotypes Additionally, exploring the interplay between tolerance and host immune responses may provide further insight into how AFDT contributes to persistent infections. As the global burden of candidiasis rises alongside increasing drug resistance, understanding and addressing AFDT will be critical for improving therapeutic outcomes.

## Materials & Methods

### Yeast strains, media and antifungal drugs

*Nakaseomyces glabratus* wild type (typed ATCC2001-CBS2001) strain was purchased from the American Tissue Culture Collection and the clinical isolates (listed in Table 1) were obtained from Dr. Derek Fairley at Royal Victoria Hospital, UK. Strains were routinely cultured in Yeast extract Peptone Dextrose (YPD) rich agar (yeast extract: (10 g/L) Sigma Aldrich 8013-01-2; bacto peptone (20 g/L); Difco: 9295043; D(+)glucose (20 g/L) G7021:sigma-aldrich, bacto-agar: sigma A5306-250G) incubated at 37 °C and with shaking at 180 rpm (liquid cultures). Streaked plates were stored at 4 °C for up to 2 weeks. For long-term storage, YPD cultures were supplemented with 15% filter-sterilised glycerol and stored at −80 °C.

In terms of media variability, casitone media (Thermofischer: 225930) and Sabouraud dextrose broth (Sigma Aldrich S3306) media were plated onto agar plates for use. Synthetic complete (SC) media with 20% glucose (Merck Y1251 Yeast Nitrogen Base Without Amino Acids and Ammonium Sulphate (6.7 g/L), 0.22 g/L amino acid mix (L-histadine: 1 g L-tryptophan: 1 g Uracil: 1 g, L-leucine 1 g), Yeast synthetic drop out mix (-his -ura -leu and -trp); Sigma-Aldrich # Y1771) was prepared as the growth media.

For the disc-diffusion assay, the antifungal drug discs utilized were Fluconazole (FCZ) (OxoidTM CT1806B), Ketoconazole (KCZ) (Ridacom SD277-5CT) and Voriconazole (VCZ) (Ridacom sd274-5CT). For the broth-microdilution assay, the drugs Fluconazole (Biorbyt #orb1223822), Voriconazole (Biorbyt #orb134756) and Ketoconazole (Biorbyt #orb545799) purchased in powder form were utilized. The following stock solutions were prepared dimethyl-sulfoxide (DMSO): FCZ, 64 mg/ml, KCZ, 25 mg/ml and VCZ 20 mg/ml and stored at −20 °C for further use.

### Anti-fungal Minimal Inhibitory Concentration (MIC_50_) determination

CLSI M27-A2 protocols (45) for MIC quantification were followed with minor alterations. Briefly, freshly streaked colonies were grown in YPD overnight, reseeded the following morning to OD A600 ∼0.1 and grown to log phase (OD A600 ∼0.5). Cells are harvested and resuspended in sterile phospho-buffered saline (PBS) to an OD A600 0.1 (approximately 3E +06 cells). A 2-fold serial dilution for 10 concentrations of each drug namely FCZ (0.5-256 µg/ml), VCZ (0.06-32 µg/ml) and KCZ (0.25-128 µg/ml) were made following the CLSI standards with some adjustments in the concentration range. Further, instead of using the standard RPMI media (supplemented with MOPS buffer) suggested by the CLSI protocol, we have optimised the assay with SC-Complete media, whose preparation has been previously described.

In a round-bottom 96 well plate (Sarstedt), approximately 3E+04 with a total volume of 200 µl composed of media and drugs were added per well. The readings were taken at 600 nm using a FLUOstar® Omega (BMG Labtech) microplate reader. The readings were programmed to be recorded every 30 minutes with 22 flashes per well without path length correction starting from the 0 hr time point to a maximum of 48 hr time point. The program specifications were further set to 22 flashes per well without path length correction and orbital shaking at 200 rpm. The plates were placed in between the reading intervals, adhering to the temperature requirements of 37 °C. The OD values derived at the 24-hour time points from the plate reader were analysed to obtain the cut-off concentration of the drug at which there was an inhibition for an overall 50 % population of the cells over a set of three replicates.

### Fraction of Growth (FoG) Assay to measure Antifungal Drug Tolerance

FoG assays are based on modified disk diffusion assays (DDA) that were executed according to the CLSI M44-A2 standardized protocol (46) with minor adjustments according to the optimization requirements. Briefly, a freshly streaked colony from YPD agar plates was re-suspended in 1 ml sterile PBS and ODA600 adjusted to 0.5 (approximately 1.5 E+07 cells/ml. One hundred µl (∼1.5E +06 cells) were plated onto ypd plates using four to five sterile glass beads (mm diameter: Sigma #1040160500). Designated drug disks (Fluconazole 25 µg; Ketoconazole 50 µg; Voriconazole 1 µg) were carefully placed at the centre of each plate with a sterile tweezer and the plate was incubated at the required temperature (typically 37 °C). For a given strain and condition, typically three biological replicates (unique colonies) were assayed. Plates were imaged under LED illumination using an iPhone 13. The images were adjusted with to increased contrast background using Adobe Photoshop V24 according to the program specifications for better recognition of the pixel densities. Once run in the R console using the package <u>vignette</u> (41) which implements image J, the disk image program gives a set of values that includes a fraction of growth (FOG_20_, FOG_50_, FOG_80_) each denoting the number of colonies in the area where there are 20%, 50% and 80% inhibition rates. It further gives a RAD diameter (RAD_20_, RAD_50_, RAD_80_) representing the distance from the disc centre where 20%, 50% and 80% inhibition zones lie. Ideally, for our tolerance estimation, the values RAD_20_ and FOG_20_ were primarily considered following previous research studies (34). Each data set was replicated for three successive biological and technical replicas and average FOG_20_ values along with the standard error were determined and plotted using GraphPad prism V 10.2.2.

### Supra-MIC Growth (SMG) Assays

SMG assays are set up in an identical way as for MIC determination detailed above. Following MIC determination at 24 hours, cultures are incubated for a further 24 hours under the same conditions and OD A600 is recorded. To quantify SMG, the absorbance readings in wells that contain drug concentrations above the calculated MIC50 for that strain are averaged, and normalized to growth after 48 hours in the absence of anti-fungal according to this equation(33):

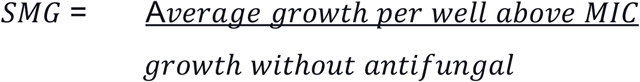

### In-vivo Galleria mellonella Nakaseomyces glabratus infection model

Last-instar larvae of *Galleria mellonella* were sourced from UK Waxworms Ltd Sheffield, UK). The experiments were undertaken following the pre-established protocols for fungal virulence testing ((38, 42, 43). Briefly, larvae were selected based on weight ranging between 250-350 mg and devoid of any visible signs of melanization(43). *N. glabratus* log phase cultures (OD A600 0.5-0.7) were harvested and cells were resuspended in sterile PBS to an OD A600 ∼16.6, corresponding to approximately 5 E+08 cells/ml. For each experiment, 20 larvae were infected with 10 µl of the yeast suspension through their injection into the rear left pro-leg using a Microliter™ syringe (Hamilton) with a 30-gauge needle (BD Microlance™ 3) of 0.5 inches. Control groups underwent similar procedures, receiving either 10 µl of sterile PBS or a needle stab without injection. After 30 minutes, fluconazole or placebo was administered to each group of larvae by a second 10 µl injection into the rear right pro-leg. The fluconazole was injected with a dosage of 12 mg/kg, while control samples were given a second injection of PBS containing an equivalent volume of DMSO. Larvae were incubated in standard 60 mm Petri dishes at 37 °C and in the dark and monitored every 24 hours for 7 days to assess viability. Larval mortality was determined when they ceased movement in response to touch and subsequent observations for cocooning, death and survival were recorded. 3 sets of experimental replicates were conducted independently, and the data was combined to create Kaplan-Meier survival curves. Statistical significance between experimental conditions was assessed through pairwise comparisons using the Log-Rank (Mantel-Cox) test.

## Acknowledgements

This work was supported by funding from the Department for the Economy (DfE), Northern Ireland, United Kingdom. We would like to thank Dr. Derek Fairley (Royal Victoria Hospital) for providing the clinical isolates. We are also grateful to Dr. Aleeza Gerstein for her valuable guidance in optimising the FoG estimation. Finally, we acknowledge the School of Biological Sciences, Queen’s University Belfast, for providing the necessary facilities and support.

## Author contributions

Dr. Edel Hyland and Sreyashi Acharjee contributed equally to this work. Both authors were involved in the conceptualisation, design, execution, and analysis of the study. They jointly wrote and revised the manuscript. All authors have read and approved the final version of the manuscript.

Declaration of generative AI and AI-assisted technologies in the writing process During the preparation of this work the author(s) used ChatGPT in order to improve readability and decrease word count. After using this tool/service, the author(s) reviewed and edited the content as needed and take(s) full responsibility for the content of the publication.

## Supplementary figures

**Figure S1:**
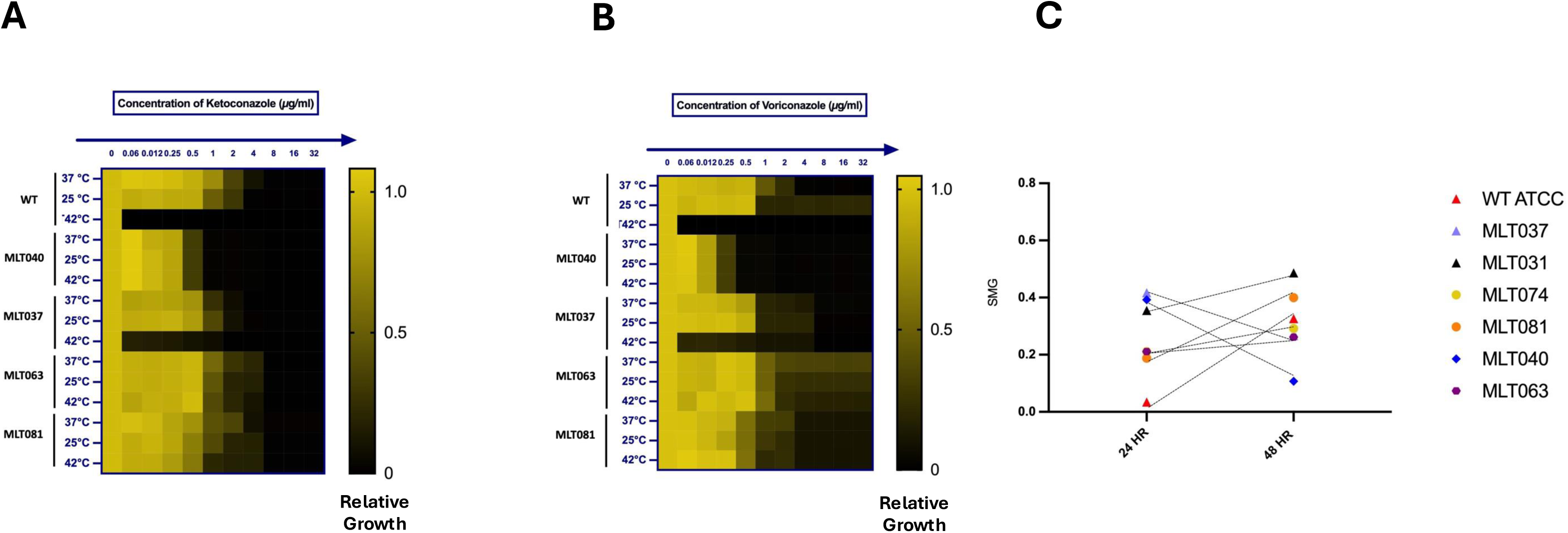
Clinical isolates of *N. glabrata* exhibit varying levels of drug tolerance under different temperature conditions. Panels A and B show SMG (Supra-MIC Growth) values for clinical isolates, calculated as described in Figure 3. These experiments were conducted in SC-complete yeast media at three different temperatures: 25 °C, 37 °C, and 42 °C, using two-fold increasing concentrations of antifungal drugs for the following drugs (A) ketoconazole(B) voriconazole. (C) SMG values as a measure of growth in the presence of fluconazole (FCZ), specifically at supra-MIC drug concentrations. Growth was assessed at both 24-hour and 48-hour time points to capture temporal dynamics in tolerance.

